# Heterogeneous distribution of mRNAs within flight muscle fibers, and implications for function

**DOI:** 10.1101/2020.09.07.286260

**Authors:** Aditya Parekh, Kunal Chakraborty, Devam J Purohit, Shaik Naseer Pasha, R. Sowdhamini, K. VijayRaghavan, Dhananjay Chaturvedi

## Abstract

Muscle heterogeneity has been explored in terms of fiber-type distribution, structural organisation, and differences at their junctions with neurons and tendons. We amplify on such observation to additionally suggest that muscle syncytia have nonuniform protein requirements along their length, deployed for developmental and functional uses. An exploration of regionalized proteins or their mRNA across muscle syncytia has not been done. We investigated mRNA localization in regions of *Drosophila melanogaster* dorsal longitudinal muscle (DLM) syncytia over their entire transcriptome. Dissection of muscle regions, their RNA-seq and stringent Differential Gene Expression analysis indeed reveals statistically significant regionalization of nearly a hundred mRNA over the length of DLMs. Functions of over half of these genes require experimental verification. A preponderance of mRNA coding for catabolic and proteolytic enzymes is conspicuous among transcripts enriched in the posterior of DLMs. Our findings provide a foundation for exploring molecular processes that contribute to syncytial maturation and muscle homeostasis in a spatially non-homogenous manner.

## Introduction

Specialized mononucleate cell types can limit protein function in time and to subcellular regions by regulating the availability of corresponding mRNA[1]. Transcript localization to subcellular regions has been shown in several mononucleate contexts such as embryos and neurons in many organisms[2,3]. A single cell with multiple nuclei constitutes a syncytium. Regulating protein availability in subcellular regions of syncytia can involve the use of a variety of regulatory mechanisms. Internal parameters such as organellar arrangement, cytoplasmic viscosity and cellular dimensions are involved in protein localization in ways that are sometimes not seen in mononucleate cells. mRNA localization and its regulatory consequence in syncytia have been best examined in the Drosophila blastoderm, where transcript regionalization and dynamics have been linked deeply with function[4–6].

The cartography of transcripts in muscle syncytia remains inadequately explored, in contrast. Skeletal muscles are of particular interest owing to their numerous constituent nuclei, dense construction, dynamic mechanical function and large size. For specific genes, myosyncytial mRNA localization is controlled by locally limited transcription ostensibly in response to local requirements. Studies show enrichment of mRNA at the site of the function of their corresponding proteins; such as those of Talin, Myosin Heavy Chain at the myotendinous junctions[7,8] and Acetylcholine receptor-alpha subunit and Acetylcholine esterase at neuromuscular junctions[9,10]. Heterogenous expression of proteins within the same muscle syncytium has also been previously reported[11,12]. These findings support the myonuclear domain hypothesis, which suggests that individual nuclei satisfy the transcriptional requirements in the proximal cytoplasm[13]. However, it is apparent from studies on the Drosophila syncytial blastoderm that regionally localized RNA and proteins can exert long-range effects, and not just serve local requirements. This hypothesis has not been tested in muscles, in no small measure due to the unavailability of information on transcript distribution across the fiber.

To exhaustively characterize heterogeneity at the level of mRNA localization over the whole transcriptome in syncytia, we chose the adult Drosophila dorsal longitudinal muscles (DLMs) for study. The DLMs are a subset of the indirect flight muscles[14] that power flight. They constitute one contractile unit, and therefore one muscle, of one fiber-type with six fibers per hemithorax[14,15]. Flight muscles constitute 65% of the body weight of adult Drosophila[16] and the DLMs are their largest sub-group. The DLMs are finely positioned within the thorax along with the other group of indirect flight muscles, the dorso-ventral muscles (DVMs). The large jump muscle and the direct flight muscles are the other muscles in the mesothorax[17,18]. The thorax also accommodates the ventral nerve chord and sections of the gut and heart[19]. From this tight arrangement, we specifically dissected out DLMs arbitrarily compartmentalized into anterior, medial & posterior from thoraces and sequenced their mRNA. Thus, our experiments are aimed to detect transcript heterogeneity along the axis of what is essentially, anatomically, a structure with little variation.

Our internally consistent and stringently filtered data clearly indicate enrichment of small subsets of mRNA in each muscle region. We hypothesize that several of these localized mRNA and the proteins they encode could play roles not just in muscle fiber function, but also in short- and long-range regulation of growth and homeostasis. The microscopic examination of the Drosophila nuclear blastoderm stage did not reveal major intracellular heterogeneity which was revealed by later genetic and molecular tools[6,20]. Similarly, we suggest that the mRNA heterogeneity we observe in these uniform and stereotypic muscle[21] will reveal regulatory roles in both growth and repair.

## Results

### Workflow to identify heterogeneity in mRNA localization in dorsal longitudinal muscle

The significant component of Drosophila thorax is the flight muscles of which a subset is the Dorsal Longitudinal Muscles (DLMs). Drosophila Dorsal Longitudinal Muscles (DLMs) are a part of adult indirect flight muscles and these syncytia run anterior to posterior (of Drosophila) on either side of the thoracic medio-sagittal plane (Fig 1C, D). The sagittal view of the DLMs gives us a picture of stereotypic linear arrangements of nuclei and innervations of motorneurons coming from the ventral nerve cord, forming the essential system for neuromuscular coordination (Fig 1C, D). DLMs are attached to the body wall through tendons on either side[22,23] (Fig 1C) and are extensively tracheated[24]. The schematic of the transverse section of the thorax displays six DLM fibers arranged one beneath the other and guarded by DVMs (Dorsoventral muscles) and TDT (tergal depressor of the trochanter) (Fig 1A). Fig 1B shows thorax dissected in transverse section (cut on the two sides of the wing hinges). Six pairs of DLMs are visible on either side of the mid-line (marked by star), which are guarded by DVMs (another subset of indirect flight muscles, marked by circle) and TDT/jump muscle (marked by triangle) at the sides (Fig 1B). We speculated that differences in the proximity to tendons, innervation and tracheation, apart from hitherto unannotated internal cytological differences along the length of DLMs (~500-800μm long) may contribute to a possible difference in mRNA localization[22,25]. In the absence of obvious landmarks separating DLMs into lengthwise sections, we chose to define regions by cross-sectional planes separating the three pairs of legs. In this way, we defined Anterior (A), Medial (M) and Posterior (P) regions in these syncytia (Fig 1D). Assessment of differential mRNA localization in these DLM regions required their dissection along with a non-muscle control tissue, isolation of RNA and analysis of differences in transcript levels. For our study, we chose gonads from the same animals as a non-muscle control tissue.

**Fig 1.**
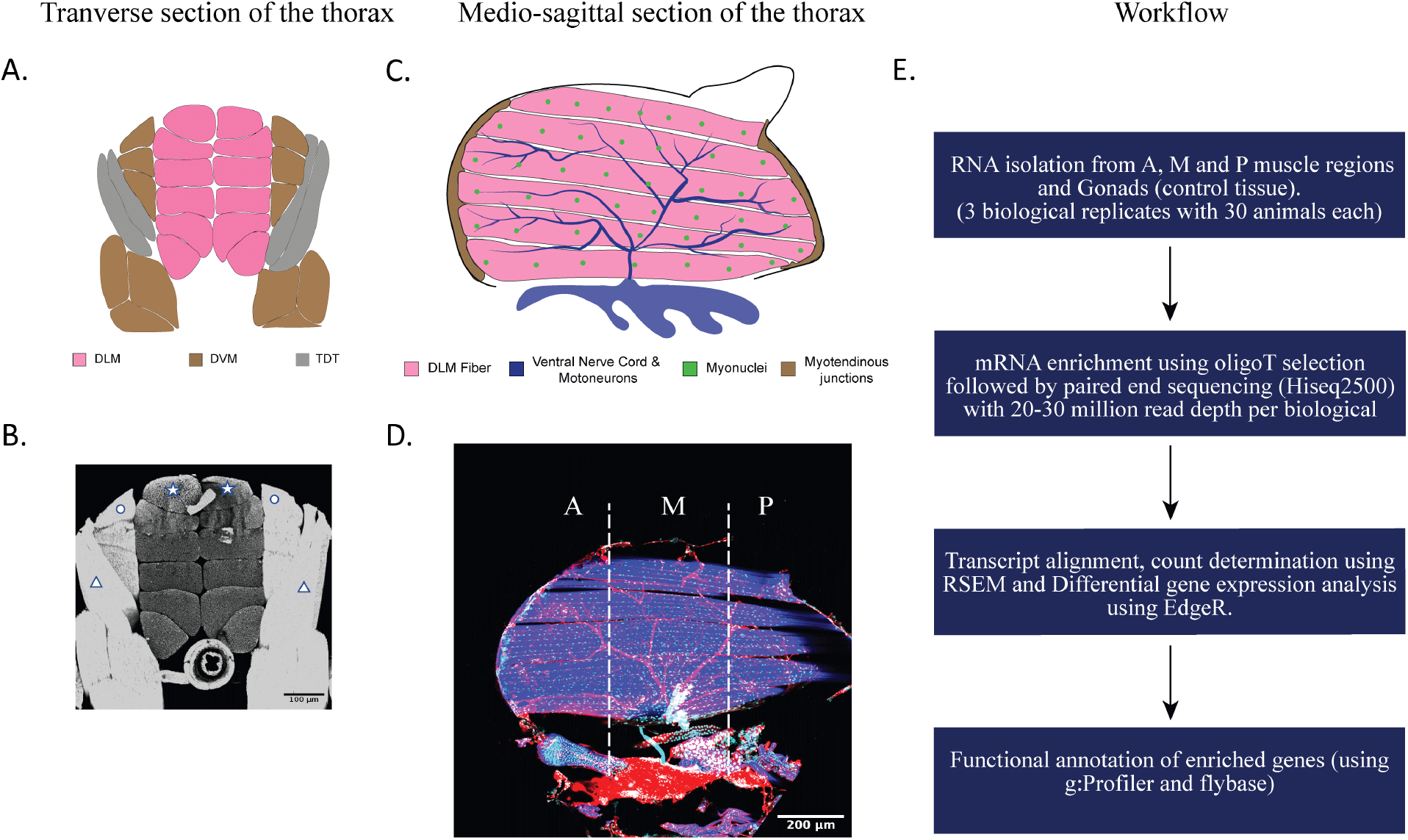
Work-flow for heterogeneous mRNA localization analysis in DLMs. **A.** Schematic representation of the transverse section of the Drosophila across the medial region. Six pairs of DLMs (Dorsal Longitudinal muscle), arranged one after the other across the midline is shown in pink. This is flanked by DVM (Dorsoventral muscle) (brown) and TDT (tergal depressor of trochanter) (grey) on both sides of the median thorax. **B.** Transverse section of the thorax (cut on the two sides of the wing hinges) showing arrangement of 6 pairs of DLMs arranged across the midline. First DLM fiber on each side is marked with a star. Other 10 unmarked DLM fibers are arranged beneath them. DLM is guarded by another type of indirect flight muscle, DVMs, on both the sides (marked with a circle). TDT/jump muscles at the sides are also shown (marked with a triangle). The image shows labelling of actin by phalloidin. **C.** Schematic representation of the sagittal section of Thorax, dissected across the midline. Six pairs of multinucleated DLM fibers run anterior to posterior (left to right, demonstrated in pink). Collinear myonuclei are shown in green. DLMs are attached on both the sides to tendons via myo-tendinous junctions (brown). Motor neuron (blue) branch from the ventral nerve cord (blue) and form junctions with the muscle fibers. **D.** Medio-saggital section of the thorax displays DLMs in the hemithorax with phalloidin (blue), syncytial nuclei (cyan) innervated motor neuron and ventral nerve cord (in red). DLMs run anterior to posterior of Drosophila (from left to right) on either side of the medio-sagittal plane. The DLM is divided into three regions Anterior (A), Medial (M) and Posterior (P) by the dotted line separating the three pairs of legs. These lines were further used to make an orthogonal cut (w.r.t the anterior-posterior axis) in the whole thorax for dissecting out A, M, P regions of DLM. **E.** The workflow starting from RNA isolation to analysis is demonstrated. Total RNA was isolated from the A, M, P and Gonads (G). Tissues from a total of 30 flies constituted one biological replicate which was further divided into two technical replicates during the sequencing run. 3 biological replicates for each sample type (A, M, P, and G) constituted the whole study. Subsequently, mRNA was isolated by oligodT selection and paired-end sequencing was performed using Hiseq2500 for each sample including technical replicates. Sequence depth was maintained between 20-30 million reads for each biological replicate. After sequencing, transcripts were aligned and counted using RSEM tool. edgeR was used to perform Differential Gene Expression and the functional annotation of the enriched genes was done with the help g:Profiler tool and manual search via FlyBase.

The workflow to achieve this goal is described in Fig 1E. Muscle regions were dissected out in the following manner as described in materials and methods. Briefly, each 1day post-eclosion (p.e.) Drosophila was submerged in 100% ethanol. Ethanol rapidly dehydrates tissue while preserving RNA *in situ*. Upon removal of the head of these ethanol submerged animals, internal thoracic tissue became considerably rigid due to dehydration. The remaining intact thorax with the legs and abdomen attached, was immobilized ventral side up on a glass slide and dipped in liquid nitrogen till frozen solid. Before the tissue could thaw, two cuts in the entire cross-section between the three sequential pairs of legs separated the thorax, including DLMs, into desired A, M and P regions. Owing to their size and distinct appearance, DLMs could be picked out with fine forceps without disturbing adjacent muscles and stored in 100% ethanol on ice. Gonads were isolated from the same animal. For statistical confidence in our results, tissue was collected in 3 sets (biological replicates) from 30 animals per set. After RNA was isolated from each muscle regions and gonads separately, it was subjected to quality check, sequencing, and analysis as described in the workflow, described in Fig 1E (also materials and methods 2 and 3).

**Fig S1.**
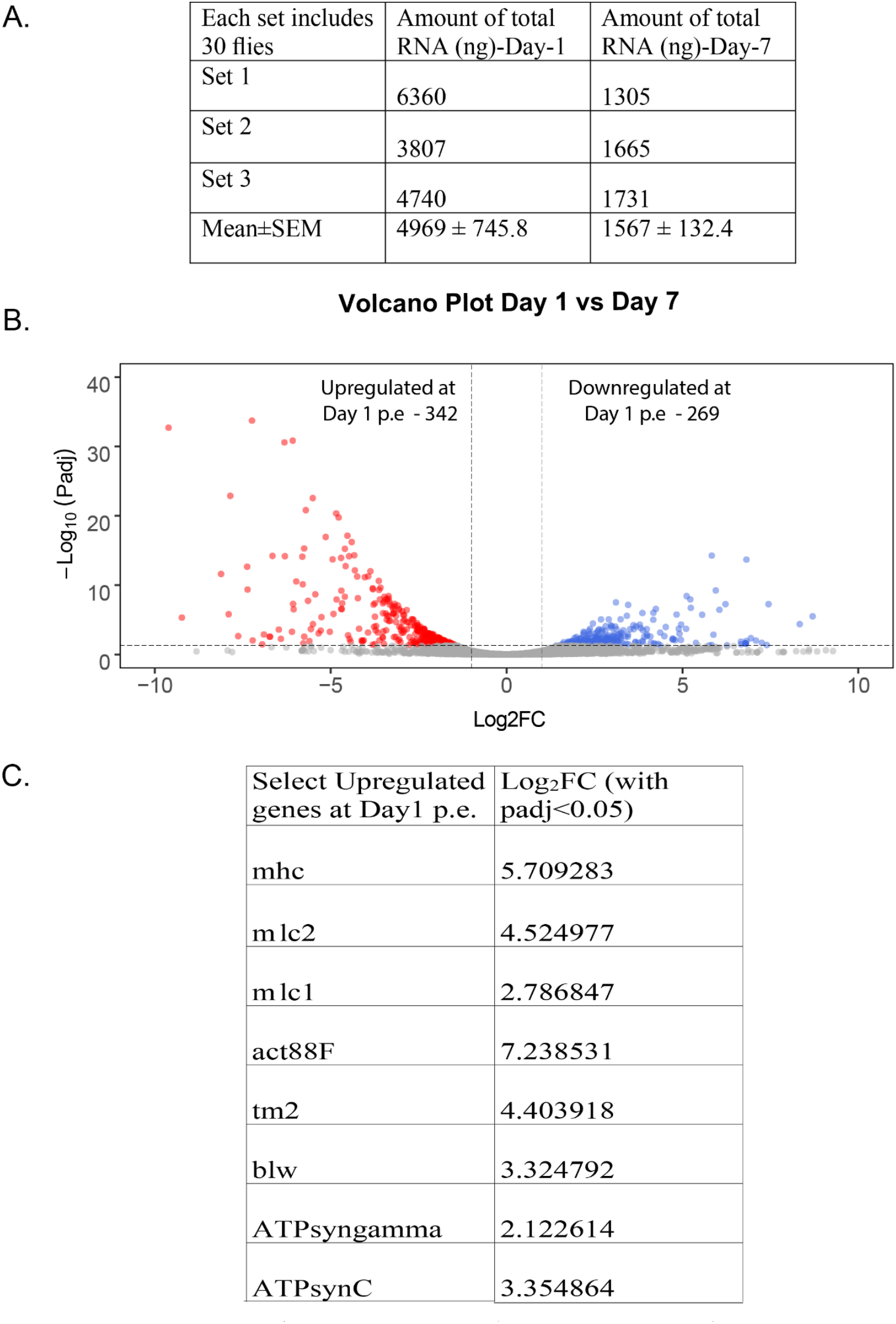
Transcriptome analysis of Day-1 and Day-7 p.e. DLMs. **A.** Comparison of total RNA isolated (ng) from all six DLMs of 90 animals divided into three sets, between Day-1 p.e. and Day-7 p.e.. The Mean of total RNA is shown along with Standard error of mean (SEM). **B.** Volcano plot showing differentially regulated genes between Day-1p.e. and Day-7 p.e. DLMs with log_2_ (fold change) > 1 and padj < 0.05. Red dots and blue dots represent the number of genes up-regulated (342) and down-regulated (269) at Day-1. Each dot represents one gene. **C.** List of select genes (8 genes) out of top 100 genes (based on average normalized counts/base mean values) upregulated at Day-1p.e. with respective log_2_ (fold change) values (Padj < 0.05)

We ascertained that total RNA isolated from ninety Day-1 p.e. DLMs in three different sets (30 animals in each set) was 3.2 times the amount isolated from the same number of Day-7 p.e. DLMs (Fig S1A). These RNA isolates also include non-coding RNA and ribosomal RNA. Simultaneously, comparison of transcript levels between Day-1 p.e. DLMs in this study with previously published [22] Day-1 IFMs (of which DLMs are an integral part) was performed (materials and methods 5). Analysis revealed that close to 22 percent genes were significantly upregulated or downregulated (Log2FC>1; Padj<0.05) (Supplemental table 1) between the two datasets. This is probably due to the difference in tissue enrichment and dissection methodology used for RNA Sequencing between this study and previously published data [22]. Differential Gene Expression analysis (supplemental file 1) between Day-1 and Day-7 p.e. from the read count values (supplemental file 2) demonstrate fold-change regulation of genes with their padj values (materials and methods 6). 342 genes are downregulated at Day-7 p.e. while 269 genes are upregulated in the course of this duration of development/ageing, as illustrated in the volcano plot (Fig S1B, supplemental file 1, sheet2). Among the top 100 enriched genes (based on average normalized counts or base mean), 44 genes were upregulated at Day-1 in relation to only 3 genes at Day-7 p.e. (supplemental file 3). Within those 44 genes, select genes are shown in Fig S1C along with their Log_2_FC. The large change in mRNA levels, especially of genes essential to muscle function like *act88F* and *mhc, mlc1, ATPsync* and t*m2* and others (Fig S1C) may be due to underlying growth and maturation processes, though animals are capable of flight at both times. Because of the larger amounts of total isolated RNA, significant enrichment of vital muscle specific genes, and possible local requirements of proteins in processes of maturation, we chose to proceed with Day-1 p.e. animals for further analysis.

**Supplemental table 1.**

| Differentially regulated genes (Filter: genes with “0” reads in all samples) | Upregulated | Downregulated | Total |
| --- | --- | --- | --- |
| Percentage regulation (%) | 11.2 | 10.7 | 21.9 |
| Number of genes | 1345 | 1287 | 2362 |

Subsequent to RNA isolation and sequencing, internal consistency formed the basis for choosing which datasets to proceed with for further analysis. Internal consistency of sequence data also serves to validate our technique of tissue isolation and all subsequent steps. Counts per million derived from the expected count in each dataset were used for pairwise comparison across all biological replicates (Anterior_1, Anterior_2, Anterior_3, Medial_1, Medial_2, Medial_3, Posterior_1, Posterior_2, Posterior_3, Gonads_1, Gonads_2, Gonads_3) and their technical replicates (Anterior_1T, Anterior_2T, Anterior_3T, Medial_1T, Medial_2T, Medial_3T, Posterior_1T, Posterior_2T, Posterior_3T, Gonads_1T, Gonads_2T, Gonads_3T). The proximity of these datasets to each other based on read counts for each gene is represented on a multidimensional scaling (MDS) plot. (Fig 2A, materials and methods 4). In all cases technical replicates of each region and tissue are proximate to each other, validating our technique and sequencing. Gonad datasets are significantly separated from muscle datasets showing that our method of tissue isolation prevents significant cross-contamination of transcripts. This is further illustrated by the several fold enrichment of muscle associated gene read counts (such as *mef2, act88F, mhc, unc-89* among others) in A (replicates of Anterior_2, Anterior_3), M (replicates of Medial_2, Medial_3), P (replicates of Posterior_2, Posterior_3) but not in G (replicates of Gonads_2, Gonads_3) (Fig 2B) (materials and methods 7). Conversely, several-fold enrichment of gonad associated gene read counts (such as *piwi, chic, vasa, mael* among others) is seen in G but not in A, M, P (Fig 2C) (materials and methods 7). The MDS plot (Fig 2A) reveals that datasets Anterior_2, Anterior_3 (and their technical replicates), Medial_1, Medial_2, Medial_3 (and their technical replicates), Posterior_2, Posterior_3 (and their technical replicates), Gonads_1, Gonads_2, Gonads_3 (and their technical replicates) are proximately placed within the group but sufficiently distant from each other. To maintain pairwise comparisons in read-count analysis, Anterior_1, Medial_1, Posterior_1, Gonads_1 and their technical replicates were omitted from further analysis. Statistically, these data validate the dissection and sequencing technique.

**Fig 2.**
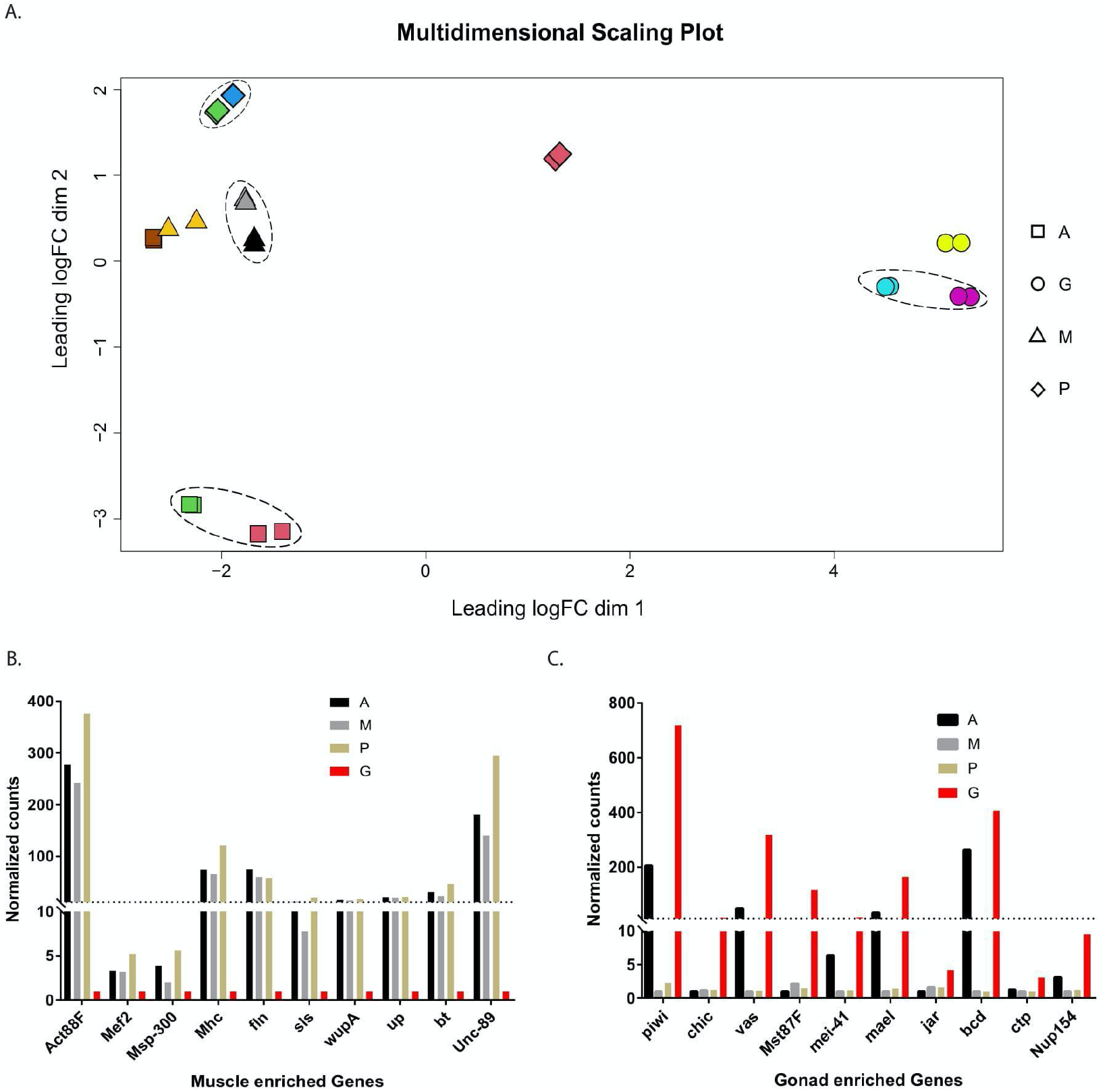
Transcriptome comparison between muscle regions and gonad datasets. **A.** A multidimensional scaling plot to represent pairwise similarity between transcriptome datasets from biological triplicates of muscle regions and gonads [Anterior (A): is represented by a square □, Medial (M) is represented by a triangle△, Posterior (P): is represented by a diamond ◊, Gonad (G): is represented by a circle ○. The technical replicates are represented by the same geometric symbol and colour. Each symbol represents a dataset. The symbols for each muscle region sample and gonad is tabulated at the right. The distance between the symbols in this cartesian space representation is proportional to the dissimilarity of corresponding datasets. Two biological replicates from each dataset were considered for further analysis and are encircled. One biological replicate dataset from each sample was excluded to maintain pairwise comparisons. **B.** Comparison of the normalized read counts for genes with known expression in muscle regions with corresponding read counts in gonads, normalized to the smallest read value across A,M,P and G (A: Black, M: Grey, P: Beige represent muscle regions; G: Red represents Gonad) **C.** Comparison of the normalized read counts for genes with known expression in gonads with corresponding normalized read counts in muscle regions normalized to the smallest read value across A,M,P and G (A: Black, M: Grey, P: Beige represent muscle regions. G: Red, represents Gonad).

**Fig 3.**
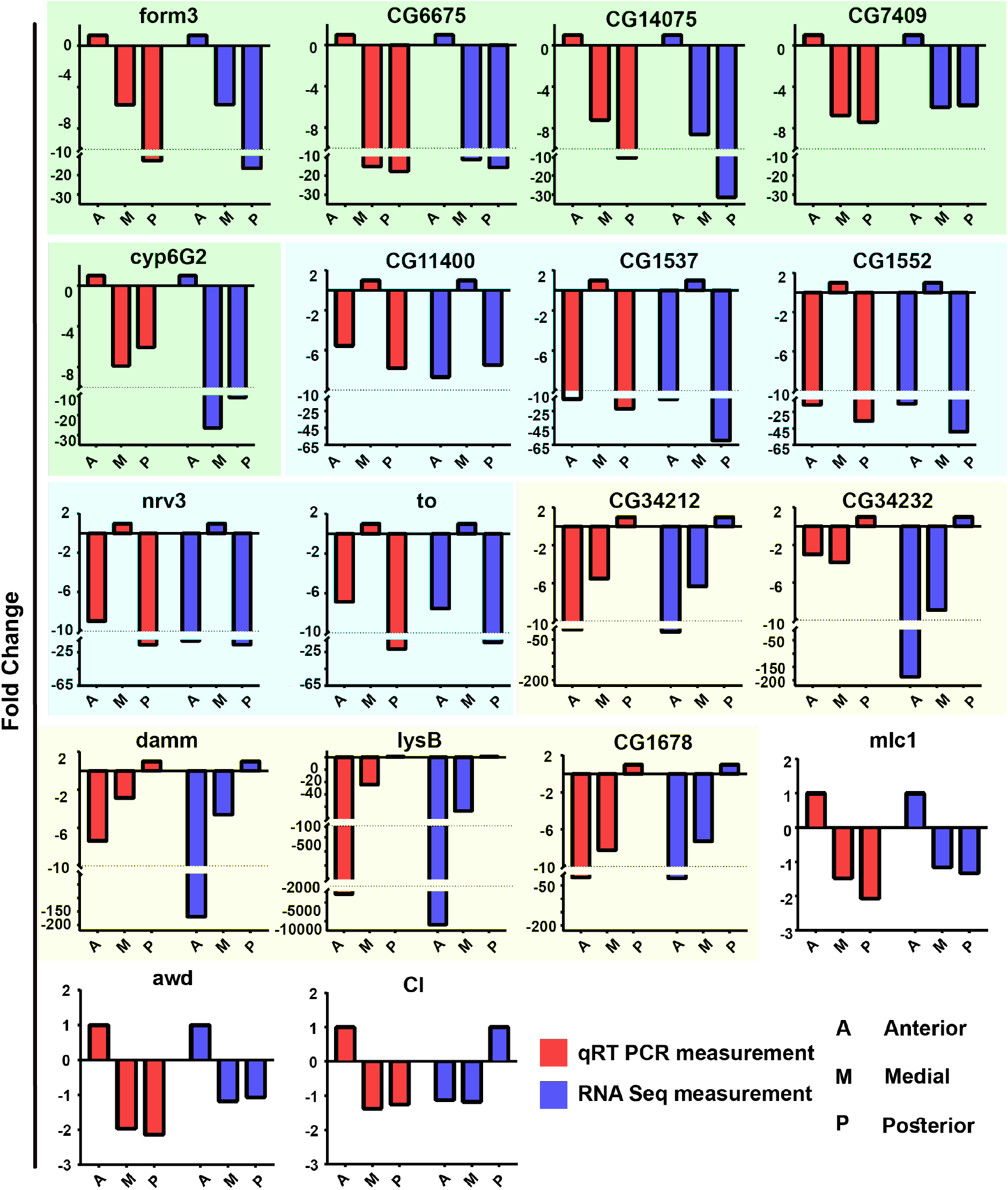
Validation of RNA-Seq read counts from each muscle region for selected genes through qRT PCR. Comparison of relative transcript levels (Fold Change: Y-axis on all plots) in different muscle regions (A,M,P, X-axis on all plots) measured through qRT PCR (red bars) and relative read counts in different muscle regions measured through RNA-Seq (blue bars). Total RNA from replicates Anterior_2, Medial_2, and Posterior_2 was used. Name of the genes are mentioned on top of each graph. mRNA of *form3*, CG6675, CG14075, CG7409 and *cyp6G2* (plotted against a green background) show enrichment in the A region; Transcripts of CG11400, CG1537, CG1552, *nrv3* and *to* (plotted against a blue background) show enrichment in the M region; Transcripts of CG34212, CG34232, *damm, lysB*, CG1678 (plotted against a yellow background) show enrichment in the P region; *mlc1, awd*, and *Cl* (in white background) showed no significant enrichment.

To be certain of enrichment of mRNA in muscle regions, RNA-seq counts would need to be validated independently through qRT PCR (material and methods 8). From differential expression analysis performed on RNA-seq data, we sampled genes that showed enrichment in any muscle regions. Fold enrichment of mRNA from these genes: *form3*, CG6675, CG14075, CG7409 and *cyp6G2* (showing enrichment in A); CG11400, CG1537, CG1552, *nrv3* (showing enrichment in M); CG34212, CG34232, *damm*, *lysB*, CG1678 (showing enrichment in P); *mlc1, awd*, and *Cl* (showing no significant enrichment) were verified from Anterior_2, Medial_2 and Posterior_2 RNA isolates through qRT PCR. Trends in fold change from the three regions and consequently enrichment, match between qRT PCR and RNA-Seq counts for most genes excluding *cl*. *mlc1*, *awd* and *cl* show little enrichment and therefore similar genes would not appear in subsequent analyses. Notably, fold enrichment of RNA-Seq counts for CG34232, *damm* and *lysB* differ by one order of magnitude from qPCR measurements, while still retaining the overall trend. Zero counts were obtained for *sgs5* and *ac* in RNA-seq data matched by little/non-specific amplification in qRT-PCR, as was evident by their Ct values (supplemental table 2). These data suggest that the sensitivity of our methods is sufficient to discern statistically significant enrichment of mRNA transcripts in any muscle region.

**Supplemental Table 2.**

| Genes | Anterior (Ct) | Medial (Ct) | Posterior (Ct) | Negative control (-RT)(Ct) |
| --- | --- | --- | --- | --- |
| *ac* | 27.90773 | 26.84555 | 26.67112 | 29.39272 |
| *Sgs5* | 32.1035 | 30.8304 | 31.33694 | 34.89787 |

### mRNA are non-uniformly localized in regions of DLMs

Having validated our methods for localized transcript assessment, we aimed to identify a candidate list of mRNA with possible biological significance. Outputs from edgeR presented us with the pairwise comparison of different muscle regions. Volcano plots (fig S2) demonstrated the number of genes which were significantly (cut-off- 1<Log2FC<-1; FDR<0.05) regulated upwards or downwards in pairwise comparison A vs M (Fig S2A), M vs P (Fig S2B), A vs P (Fig S2C). Further, complementary set of genes from pairwise comparison gave us the list of significantly enriched genes in a particular muscle region (Anterior= 432 genes, Medial=363 genes, Posterior= 256 genes), named as “unfiltered gene set” (supplemental Table 3, materials and methods 9). To derive a list of candidate genes, with likely biological relevance, we imposed a stringent filter on this output. The filter, which was named as E2F, selects for genes where the read counts across all datasets were greater than zero and enrichment of each (E) of the biological replicate of the concerned muscle region was greater than two (2) fold (F) over both the biological replicates of other two muscle regions (with FDR< 0.05)(materials and methods 10). The new shrunken list, which ensures reproduced enrichment of read counts in all regions in all our datasets, had 15 genes for Anterior, 27 genes for Medial and 52 genes for posterior (supplemental table 3) was named as “filtered gene sets”.

**Figure S2.**
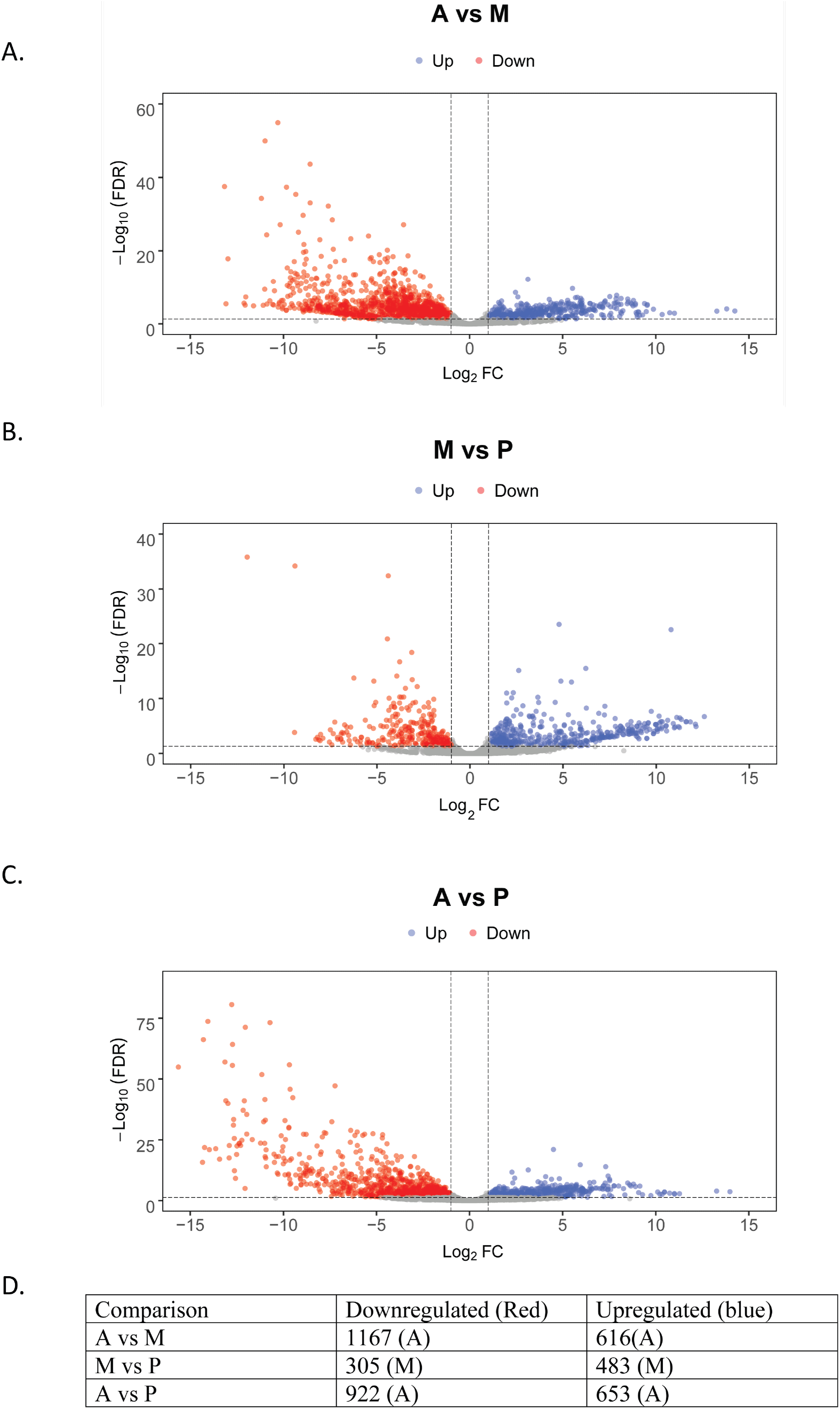
Pairwise comparison of muscle region transcriptome. Volcano plots of pairwise comparisons demonstrating significantly differentially regulated genes (FDR<0.05, Log_2_FC>1) between anterior (A) and medial (M) muscle region in A, Medial(M) and Posterior (P) muscle region in B, and Anterior(A) and Posterior (P) muscle region in C. Number of upregulated and downregulated genes in specific muscle regions derived from above volcano plots is shown in D.

**Supplemental Table 3.**

| Number of genes significantly enriched | Anterior segment | Medial segment | Posterior segment |
| --- | --- | --- | --- |
| Unfiltered gene set | 432 | 363 | 256 |
| Filtered gene set | 15 | 27 | 52 |

The levels of mRNA enrichment in various regions relative to others from “filtered gene sets” is represented in Fig 4. The Log_2_Fold Change varied from 2 to 14 in different comparisons, depicted by the gradient scale bar. To gain insight into biological functions, these genes with mRNA enrichment in various muscle regions were put through gene ontology analysis tools, g:Profiler[26]. The number of filtered candidate genes with anterior and medial enrichment was too small to fit into categories of molecular function with high significance through this analysis. Thus, manual function search (from FlyBase) was performed to understand their putative functions. Many of the anterior region genes lacked functional annotation, however, *Cyp313a1, Cyp4d21, and Cyp6g2* represented “heme binding” and “oxidoreductase” function (supplemental file 4, sheet1). For the medial region, *ara, caup* and *CG14125*, *CG34220* were interesting gene candidates in comparison as they represented “muscle cell fate commitment” and “chitin metabolism” functions respectively. The function of other medial enriched genes varied (supplemental file 4, sheet 2). The posterior region demonstrated enrichment of genes which was broadly related to the proteolytic function. More specifically, genes for molecular functions like “serine endopeptidase”, “serine hydrolase” (*yip7, epsilontry, jon65aiii, jon74e, jon99cii, jon65aiv, cg18180, cg18179, betatry, alphatry, cg8952, kappatry*) activity (FDR <10^-8^), and “hydrolase” activity (*lysB, lysD*) (FDR<10^6^) were over-represented along with few “catalytic genes” (supplemental file 4, sheet 3). Genes like *epsilontry, CG12374, lysb, jon99cii, lysd, betatry, alphatry, cg6295, lk* were shown to be associated with extracellular space. Together, based on stringent gene lists and gene ontology, the results demonstrated putative functions associated with different muscle regions. Simultaneously, it also provided the list of unannotated genes whose functions are not very well characterized but their genetic lines like RNAi, Gal4 and Mimic are present and accumulating (Supplemental file 4 sheet 1,2, and 4). Further, functional characterization of those genes in the context of the localization of their mRNA in muscle might be relevant for the understanding of cellular homeostasis.

**Fig 4.**
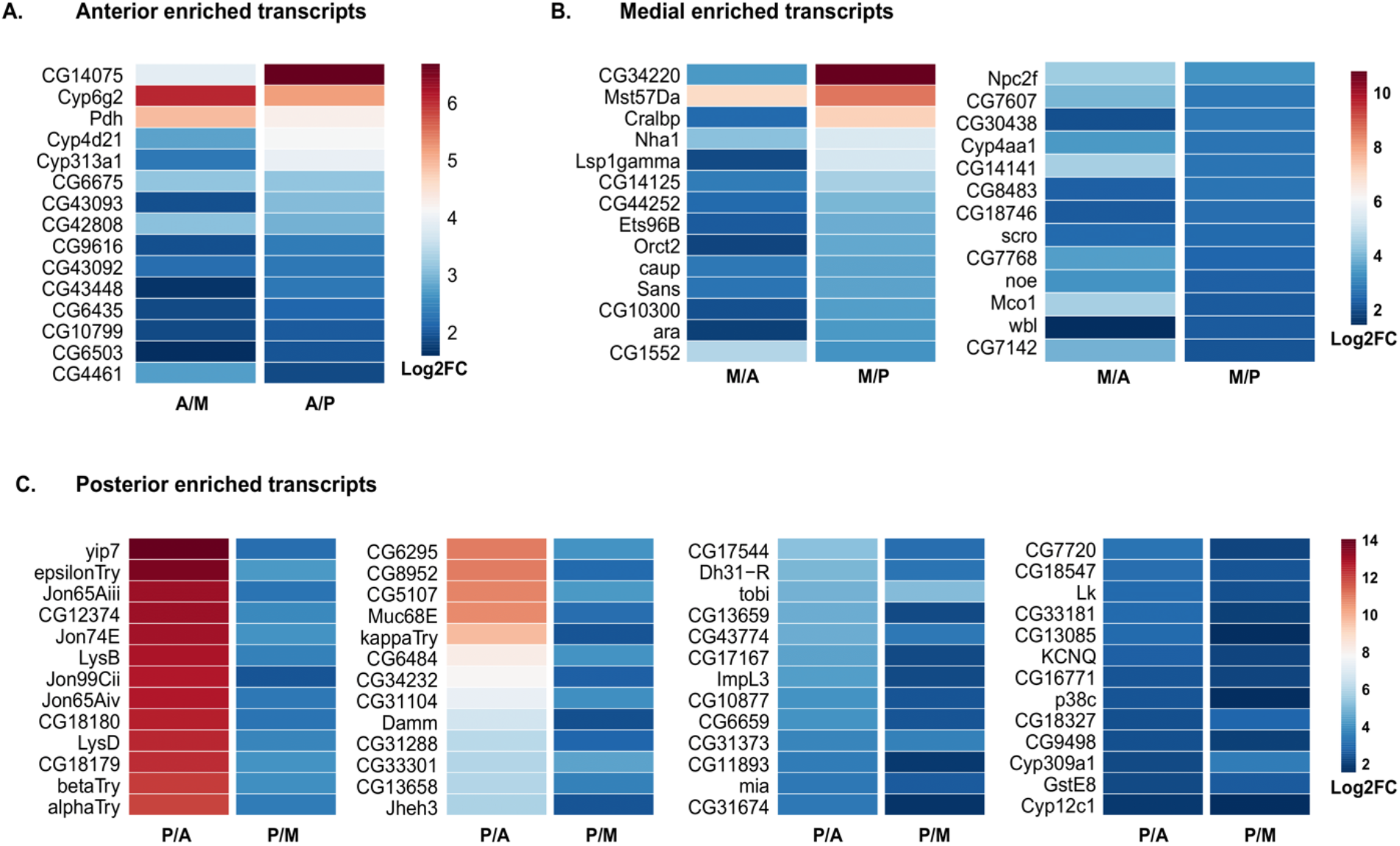
mRNA enrichment in muscle regions. Heat maps of genes from filtered gene sets showing enrichment of mRNA in different muscle regions **A.** Fold enrichments of transcripts enriched in anterior (A) relative to medial (M) and to posterior (P). **B.** Fold enrichments of transcripts enriched in M relative to A and to M. **C.** Fold enrichments of transcripts enriched in P relative to A and to M. Scales of log2 fold change shown to the right of each panel from 2(blue) to 8, 12 and 14 (deep red) in panels **A.**, **B.,** and **C.** respectively.

## Discussion

Syncytial regulation of homeostasis and growth in skeletal muscles, needed throughout an organism’s lifetime, raises interesting regulatory questions. Hypothetically, if homeostatic processes operate identically along the length of a muscle, transposing a muscle’s anterior-posterior polarity may result in a functional muscle. Mechanisms regulating the availability of active proteins within a syncytium, through possible coordination of hundreds of nuclei within a cell, in response to developmental changes in systemic energy and nutrient availability, physical cues like mechanical tension and local inputs including those from motor neurons and tendons, are an active area of study[27–30]

Protein levels and activity at any region along a syncytium may be influenced by protein transport or propagation[31], mRNA transport leading to its regionalization, ribosomal heterogeneity, localized mRNA and protein degradation, non-homogenous transcription within a syncytium or combination of these mechanisms. In this study we present a molecular analysis of mRNA localization in the DLMs in adult fruit flies. RNA-Seq analysis from lengthwise regions of the DLMs, suggests non-homogenous localization of mRNA coded for by at least 94 distinct genes (15 in anterior, 27 in medial and 52 in the posterior), some of which lack functional annotations. More sensitive methods of RNA detection may reveal a larger number of transcripts that non-homogenously localize within muscle syncytia.

At least two lines of validation of these data will be key to commence attribution of function to the heterogeneity we observe. Visually detecting mRNA, through smFISH[32] in DLMs, starting with our candidate gene list would be the first step. This is an especially relevant concern knowing that DLMs are closely and intricately associated with motor neuron arbors, tracheoles, immunocytes and tendons (in A and P regions). Though our method of RNA isolation does not deliberately exclude these tissues, the relative difference in sizes, and by extension mRNA content, likely results in an overrepresentation of muscle transcripts. Further, ascertaining protein localization, through immune-histochemistry, of these candidate geneproducts would be essential. Discrepancies between protein localization and mRNA localization will allow the regulation of protein localization in these contexts.

Revealing the contribution of non-homogenous transcript localization will require linking these to *in vivo* examination of function. Gain and loss-of localization and the consequences on function may be informative. The mRNA localization seen may be regulated by transcription from proximate nuclei. Identifying the mechanism that regulate such possibilities can now be explored by in-situ hybridisation of nuclear transcripts.

Bioinformatic tools help us categorise these gene products by annotated or putative molecular function based on homology. These gene products include enzymes such as serine proteases, lipases, oxygenases, oxidoreductases; membrane trafficking proteins, transfer/carrier proteins, proteins associated with extracellular space, ion channels and transcription factors. Perturbation studies on each of these genes, or in combination will reveal what function they play and thereby what biological processes heterogeneously operate along the length of maturing muscles. This snapshot of mRNA localization of one day post-eclosion DLMs can similarly be examined in maturing muscles and in those undergoing repair.

Published data support localized transcription in mouse muscle syncytia, through the examination of transgene expression[33]. More recently, independent single nucleus RNA-seq studies on mouse muscle syncytia[34–36] concur on heterogenous transcription in mouse muscle nuclei. Data therein strongly support previous findings and hypotheses that muscle nuclei respond to neural and tendon signals locally. The extent of neural and tendon interaction at the ends of and along the length of each muscle likely contributes to non-homogenous transcript localization.

Our findings raise an intriguing set of hypotheses: Drosophila flight muscles maybe polarized along the anterior-posterior axis as a consequence of internal long-range signaling mechanisms. The enrichment of specific proteolytic enzymes in distinct muscle regions could reflect differential myofibrillar maintenance mechanisms across the syncitium. Further, if metabolic activity could also vary across the fiber, this raises interesting new possibilities on how fiber metabolism is regulated. The exploration of this ‘Intra-syncytium regionalization’ hypothesis allows the exploration of how these large fibers function and how regulatory mechanisms so elegantly described in the Drosophila blastoderm may also apply in such cells. Given the conservation in many aspects of muscles physiology, biochemistry and function, the findings in Drosophila may have implications in other systems, including human.

Taken together, our findings provide a new approach to examine mechanisms of muscle syncytial homeostasis, growth and repair. With the development of tools for tractable intra syncytial genetic manipulation *in vivo*, much can be learnt about regionalized molecular mechanisms behind muscle homeostasis.

## Author Contributions

Conceptualization- Dhananjay, VijayRaghavan, Aditya, Sowdhamini

Experiments- Dhananjay, Aditya Parekh, Kunal, Devam

Analysis- Naseer, Devam, Aditya

Interpretation- Aditya, Dhananjay, Naseer, Devam

Writing- Dhananjay, Aditya, Kunal, Vijay

Reviewing and editing- VijayRaghavan, Sowdhamini

Funding- VijayRaghavan, Sowdhamini

## Acknowledgement

The authors would like to thank NCBS core facility, NCBS sequencing facility, NCBS server support, NCBS Fly facility. The study was conducted with the help of NCBS-TIFR Intramural funds, J.C. Bose fellowship of the Govt. of India, and ICMR RA contingency grant. Fellowship support was provided by ICMR and NCBS.

## Materials and Methods

### 1) Fly Keeping

Drosophila melanogaster *Canton S (CS)* were raised at room temperature in milk bottles on standard cornmeal agar. For accurate ageing, animals were collected within 1-hour post eclosure. Subsequently, they were distributed into 15 males and 15 females per vial and housed at room temperature. For comparison of DLM RNA levels at Day-1 and Day-7 post eclosure, vials were flipped every third day.

### 2) Dissection and RNA Isolation

For RNA isolation, all flies from one vial were transferred into one empty glass vial and placed on ice for one minute. This treatment sufficiently impeded locomotion. Flies were transferred into glass dissection dishes with 5ml of 100% ethanol. Leaving the wings intact, the head was pulled away, giving ethanol access to thoracic tissue. On a glass slide, a thin layer of glycerol was painted with a brush. Each thorax with the attached abdomen was placed individually, ventral side up and wings spread out on this glycerol patch on a glass slide, to be held in place. Held by tongs, the portion of the slide holding the thorax and abdomen was dunked in liquid nitrogen till sizzling subsided. The slide was brought out and the thorax was trisected in cross-section between the three sets of legs under a dissection scope. Sections of the thorax were distributed into wells containing 100% ethanol depending on whether they were anterior, medial or posterior muscle regions. Gonads were dissected out from the abdomen in ethanol wells and placed in 1ml ethanol tubes on ice. This dissection, ensured the dehydration of tissue and avoidance of cold shock, while preserving RNA localization and minimizing RNA contamination from other tissues. While in ethanol, DLM from each set of Anterior, Medial and Posterior regions were picked out with fine forceps and placed in respective tubes containing 1 ml ethanol on ice. The process of tissue collection from 30 animals took one hour start to finish. This constituted one biological replicate for Anterior (A), Medial (M), Posterior (P) and Gonad (G) tissue samples. Two more biological replicates were made following the same protocol.

RNA was extracted separately from DLM and gonad samples with TRIzol (Sigma) according to the manufacturer’s instructions. The total RNA from each sample, which was dissolved in DNase and RNase free water, was split into two equal parts. These aliquots were the technical replicates of the respective sample. RNA was estimated from each biological replicate and its technical replicate separately through the Qubit (Thermo Fisher Scientific) protocol. Further, the quality of RNA was verified using Bioanalyzer 2100 before proceeding for RNA-Sequencing. Equal amount of total RNA from each tube was taken forward for sequencing. In this manner, RNA-Sequencing information for the three DLM segments (A, M, P) and Gonads (G) was obtained in biological triplicates (Anterior_1; Anterior_2; Anterior_3; …) with technical replicates for each (Anterior_1, Anterior_1T; Anterior_2, Anterior_2T; Anterior_3, Anterior_3T; …). RNA of RIN 8-10 was used for subsequent analysis. mRNA enrichment was done using E7490L-NEBNext Poly(A) kit. cDNA libraries were prepared using the E7765L mRNA kit (NEB next). Paired-end read sequencing was performed using the HiSeq2500 instrument. During sequencing, read depth per biological replicate varied between 20-30 million. Library preparation and sequencing was performed by the NCBS sequencing core facility.

### 3) Read processing, transcript quantification and differential expression analysis for A (anterior), M (medial), P (posterior) muscle regions

The RNA-Seq read was assessed using FastQC and low-quality reads and adapters were trimmed using TrimGlamore v 0.4.3.1 for paired-end reads (R1 and R2), sites with PHRED scores lower than 25 and reads below 20 bp in length were removed. The RSEM v1.2.25 was run with default settings using Bowtie2 v2.3.3.1 aligner to quantify transcript expression. The paired-end reads were aligned to the Drosophila reference genome downloaded from the ensembl genome database and the corresponding GTF file (https://www.ensembl.org/info/data/ftp/index.html). RSEM computes abundance in genes based on the Expectation-Maximization (EM) algorithm. The read count obtained was filtered on the basis count-per-million (CPM) to account for differences in library size between samples to exclude genes with low counts across libraries. The only genes with a CPM >= 1 in at least two samples were retained for downstream analysis. Subsequently, for expected read count tables of genes were created for each dataset which was piped into Differential Gene Expression analysis edgeR Bioconductor package (http://bioconductor.org/packages/release/bioc/html/edgeR.html) with weighted trimmed mean of M-values normalization method and likelihood ratio tests of the glmLRT function was used to identify Differentially expressed genes. A cut-off of log_2_FC of 1 and FDR (False discovery rate) < 0.05 was used for further analysis. Three Differential Gene Expression sheets along with list of significantly regulated genes were prepared during transcriptome comparison of muscle regions, for A vs P (supplemental file 5, sheet 1,2), A vs M (supplemental file 6, sheet 1,2), and M vs P (supplemental file 7, sheet 1,2). Volcano Plot using the same cut-off was made using R package “EnhancedVolcano” by using gene ID (FBgn), log_2_FC and FDR columns from edgeR Differential Gene Expression file to display significantly regulated genes between the pairwise comparison of A vs M, A vs P, and M vs P.

### 4) MDS plot

Multidimensional Scaling (MDS) plot was used to assess the similarity between different datasets considering their read values. All the muscle (A, M, P) and the Gonad samples including their biological and technical replicates as represented in RSEM count file was used as an input. The raw reads (from RSEM) were converted to log_2_-transformed counts per million (log_2_-cpm) values using “cpm” function in edgeR, where no additional filter was used. The resulting values were then normalized using “calcNormFactors” function in edgeR. The normalized values were used to make MDS plot using plot MDS function in edgeR.

### 5) Comparing our Day-1p.e DLM transcriptome with Day-1p.e. IFM transcriptome (Spletter et al., 2018)

The raw count file from the sample sets (our Day-1p.e. DLM and Spletter et al., 2018 IFMs 1day adult) was used for Differential Gene Expression analysis using Deseq2[37] in R, which is the same as used in the Spletter et al., 2018 manuscript. “DESEqDataSetFromMatrix” function within the DESeq2 package was used in this case. The input count data in Deseq2 was filtered in R by excluding those genes which had zero reads for all samples. Cut-off of Log2Fold change>1 and Padj<0.05 was used to calculate the number of significantly regulated genes between them. Differential Gene Expression analysis and read counts are provided in the supplemental file 8 and 9 respectively.

*Two out of three biological replicates were used for Day-1 DLM sample as suggested by the MDS plot (method 4)

### 6) Transcriptome analysis of Day-1 p.e and Day-7 p.e. DLM

Both Day-1 and Day-7 p.e. condition consisted of three biological replicates. Each replicate had RNA from a total of 30 flies (15 males and 15 females). After sequencing, the quality of the RNA-Seq reads and read trimming was done using the following method as explained earlier (method 3). RSEM was used to compute the read counts (method3). The reads were further filtered and the genes which had “0” counts in all the samples including replicates were not considered for downstream analysis. MDS plot was used to assess the similarity between datasets and subsequently samples Day-7_1, Day-7_3, Day-1_1, Day-1_3 (Figure S3) were chosen for Differential Gene Expression analysis. For the Differential Gene Expression, Deseq2[37] package in R was used to prepare the list of differentially regulated genes stating the Log_2_FoldChange and Padj values. With the cut-off of Log_2_Fold change>1 and Padj<0.05, volcano Plot was made using R package “EnhancedVolcano” by using gene ID (FBgn), log_2_FC and padj columns from DESeq2 Differential Gene Expression file. Further, from Differential Gene Expression list, based on base mean value, top 100 genes were sorted. The number of significantly regulated genes within the top 100 (as per the base mean values) was used to prepare another list stating their upward or downward regulation at Day-1.

### 7) Comparison of transcripts of muscle regions versus Gonad

The expected count file (supplemental file 10_sheet1) generated from RSEM was used for depth (read) normalization. The mean of technical replicates from the expected count values was shown in supplemental file 10_sheet2. Sequencing reads were normalized for all samples with respect to the sample (G1), which had the highest total read count. For normalization, total G1 read was divided by the total read of other samples to generate normalization factor for each sample including the biological replicates. Individual read values (of all the 15682 genes) for each sample was multiplied to the respective normalization factor to generate a “normalized count file” (supplemental file 10_sheet3). Further, ten specific genes each for muscle and gonad were chosen for comparison of expression levels in the muscle (A,M,P regions) and gonad sample. The mean of the normalized read counts for ten genes were further normalized with respect to the read value of the sample which had the lowest count (supplemental file 11). Later, reads were plotted in a bar graph to compare differences in expression levels.

### 8) cDNA preparation and real-time PCR

RNA samples (stored previously at −80 degrees in RNase & DNase free water) were thawed carefully in ice. 200ng of total RNA from biological replicate Anterior_2, Medial_2 and Posterior_2 was used for cDNA preparation using SuperScript IV Reverse Transcriptase kit, Thermo Fisher Scientific (Cat.18090050), following its protocol. OligodT oligo was used to select for mRNAs only. Subsequently the cDNA was diluted to 45μl with RNase free water and 1μl of prepared cDNA was used for each real-time PCR (qPCR) reaction. Power SYBR^®^ Green PCR Master Mix from Applied Biosystems was used to perform qPCR with a total reaction volume of 20 μl. A “no template control” was used as a negative control which was prepared without the use of Reverse Transcriptase while preparing cDNA. 20 genes were chosen for qPCR. The primers were ordered from Sigma (0.05 μmoles and desalted) with a length varying between 90-150bps (the sequence is given in supplemental table 4). 500nM of each forward and reverse primer was used for one reaction. Each reaction was run in duplicates in a 96-well plate (Genetix-GX2196SS-FP) and was read in SDS 7500 fast Applied Biosystems machine. The reaction was run with the setting as given below.

| Stage | Holding |  | Cycling stage (40 cycles) |  |  | Melting Curve |  |  |  |
| --- | --- | --- | --- | --- | --- | --- | --- | --- | --- |
| Temperature | 50 °C | 95 °C | 95 °C | 60 °C | 72 °C | 95 °C | 60 °C | 95 °C | 60 °C |
| Time | 2min | 10min | 45sec | 30sec | 45sec | 15sec | 1min | 30sec | 15sec |

The difference in the Ct (cycle threshold) value was used to calculate the fold change. The sample with lowest Ct value for each gene was kept as one (which varies from sample to sample) and the relative expression of other genes were calculated using the 2^ (difference in Ct value). Fold change of each gene (calculated for A vs M vs P) from qPCR was compared with the fold change of RNA-Sequencing data (calculated from “normalized counts”).

**Supplemental Table 4.**
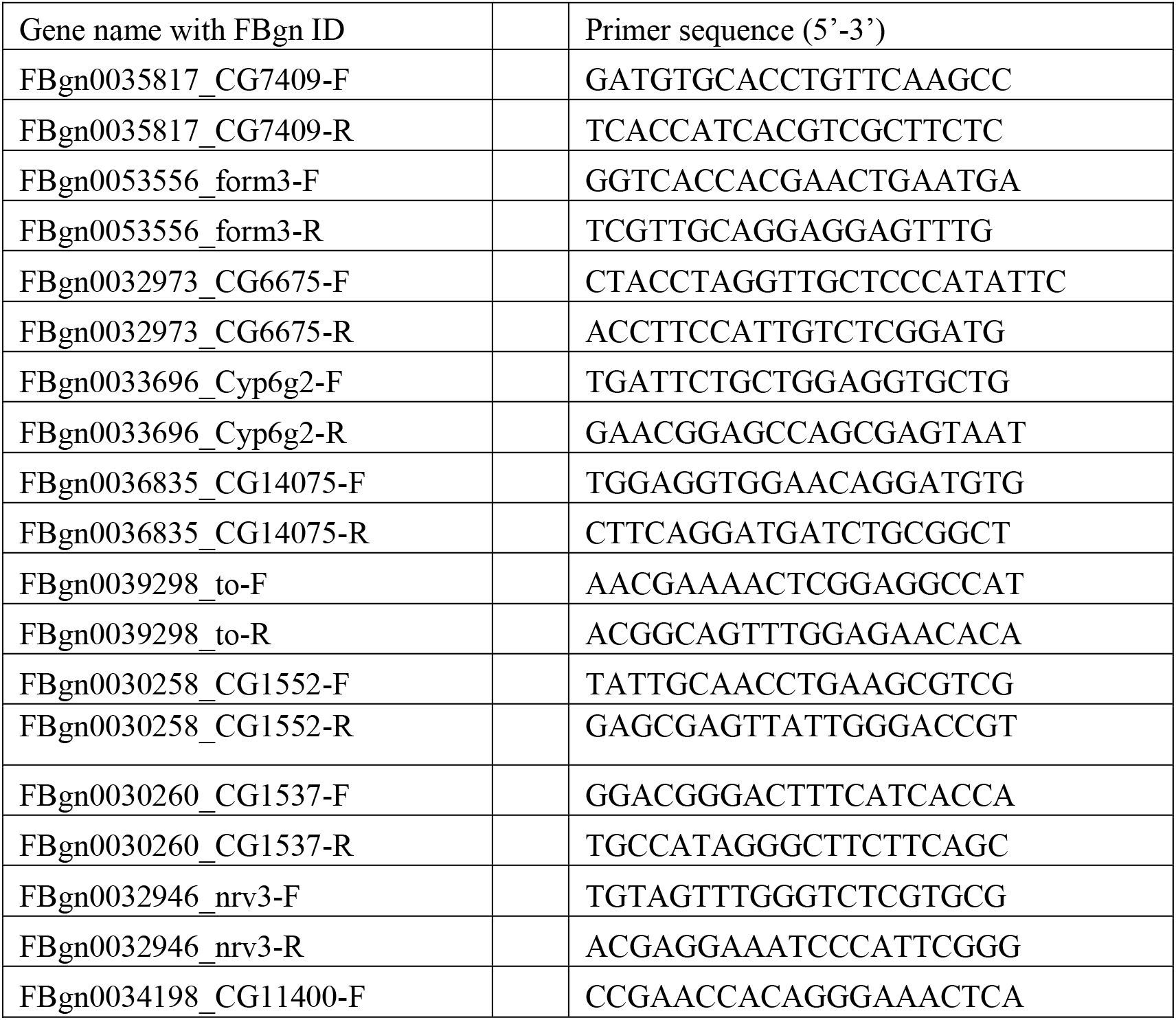
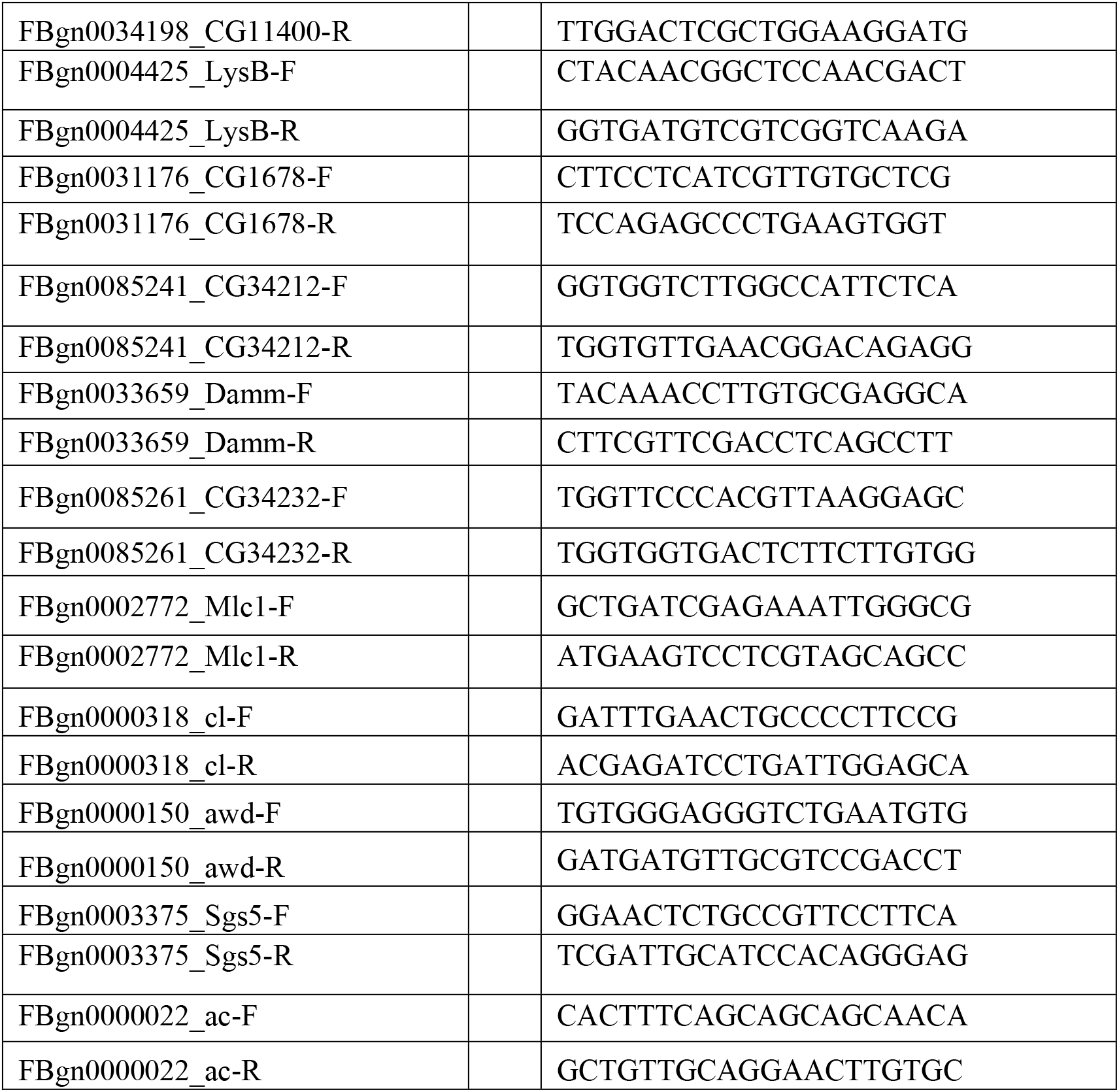

F and R denote Forward and Reverse Primer respectively.

### 9) Segment specific gene enrichment

Differential Gene Expression analysis of pairwise comparison (A vs M, M vs P, A vs P) of muscle segments yielded fold expression changes, their corresponding FDR values and other parameters. Upregulated or downregulated genes in each pairwise comparison sheet were sorted and filtered based on Log_2_FoldChange (Log_2_FC) and FDR to get the significant list of genes. The filter used was −1 ≤ Log2FC ≥1; FDR ≤ 0.05. List of complementary gene sets which were upregulated in one segment in comparison to the other two segments were identified (using pairwise comparison Differential Gene Expression sheets) and termed as “unfiltered gene sets”.

### 10) E2F filter

Since many of the genes in the unfiltered gene list showed variations within replicates, we decided to use stringent cut-off in the gene sets based on the comparison of their normalized read counts across different muscle segments. We named the filter as “E2F”. This filter selected only those genes where both the biological replicates demonstrated at-least two-fold difference with each of the two biological replicates of the other two muscle segments. Gene with “0” mean read in any of the biological replicates of any sample was not considered. This new gene list is further termed as “filtered gene sets”.

Further, a heatmap was plotted using the Log_2_FC values (calculated from Differential Gene Expression) of filtered gene sets for each of the muscle segments. The pheatmap package in “R” was used to plot the heatmap. {Raivo Kolde (2019). pheatmap: Pretty Heatmaps. R package version 1.0.12. https://CRAN.R-project.org/package=pheatmap}

### 11) Gene ontology

“Filtered gene sets” was used to identify molecular functions associated with anterior, medial and posterior muscle segments using g:Profiler tool (reference). “All genes” search parameter was selected using Drosophila melanogaster genes as a reference. Rest of the parameters were kept as default.

### 12) Immunohistochemistry and imaging

Post-eclosion two days old CS flies were taken in a glass vial and were anaesthetized by putting the vial in ice. After that immobile fly was taken in a glass well containing 1 X PBS. Head and abdomens were removed with forceps. The intact thorax was then submerged and fixed in 4% paraformaldehyde diluted in Phosphate buffered saline (1X PBS pH =7.4) for 15 minutes. Subsequently, the thorax was taken on a double-sided tape keeping the dorsal side up. With a regular razor blade, the thorax was bisected following the midline dividing the thorax into two hemi-thoraces. Hemi-thorax samples were then subjected to two washes of 0.3% PTX (PBS + 0.3% Triton-X) and 0.3% PBTX (PBS + 0.3% Triton-X + 0.1 %BSA) for 15 m each. To mark motorneurons goat anti-HRP(1:400, Jackson Immunoresearch) primary antibody staining was performed for overnight at 4°C on the shaker and secondary donkey anti-goat 568(1:400, Molecular Probes) was added following four washes of 0.3% PTX. Excess of unbound secondary antibodies was removed by three washes of 0.3% PTX. To mark actin filaments, Phalloidin 647(1:400, Invitrogen) was used by keeping on a shaker for 30 minutes at room temperature. Again, the sample was given three washes of 0.3%PTX. To mark nuclei, Hoechst 33342 (2 μg/mL, ThermoFisher) was used. Hemi-thoraces were washed thrice with 0.3% PTX and finally mounted with a mounting media of 0.3% n-propyl gallate in 90% glycerol in 1X PBS (pH = 7.4). For transverse sectioning, anaesthetized intact fly was placed on a standard glass slide with little glycerol. The whole fly was submerged in liquid nitrogen to freeze and then was followed immediately by the transverse sectioning on the two sides of the wing hinges. From fixation to mounting the protocol was the same as mentioned above and only stained with Phalloidin 568 (1:400, Invitrogen).For imaging, Olympus FV 1000 and Olympus FV 3000 confocal scanning microscope was used for image acquisition. Images were processed using Fiji software[38]

### 13. Availability of Raw sequence and metadata

Raw transcript reads are available from NCBI Short Read Sequence archive (SRA), with bio-sample accession ID, under the Bioproject accession PRJNA644535.

**Figure S3.**
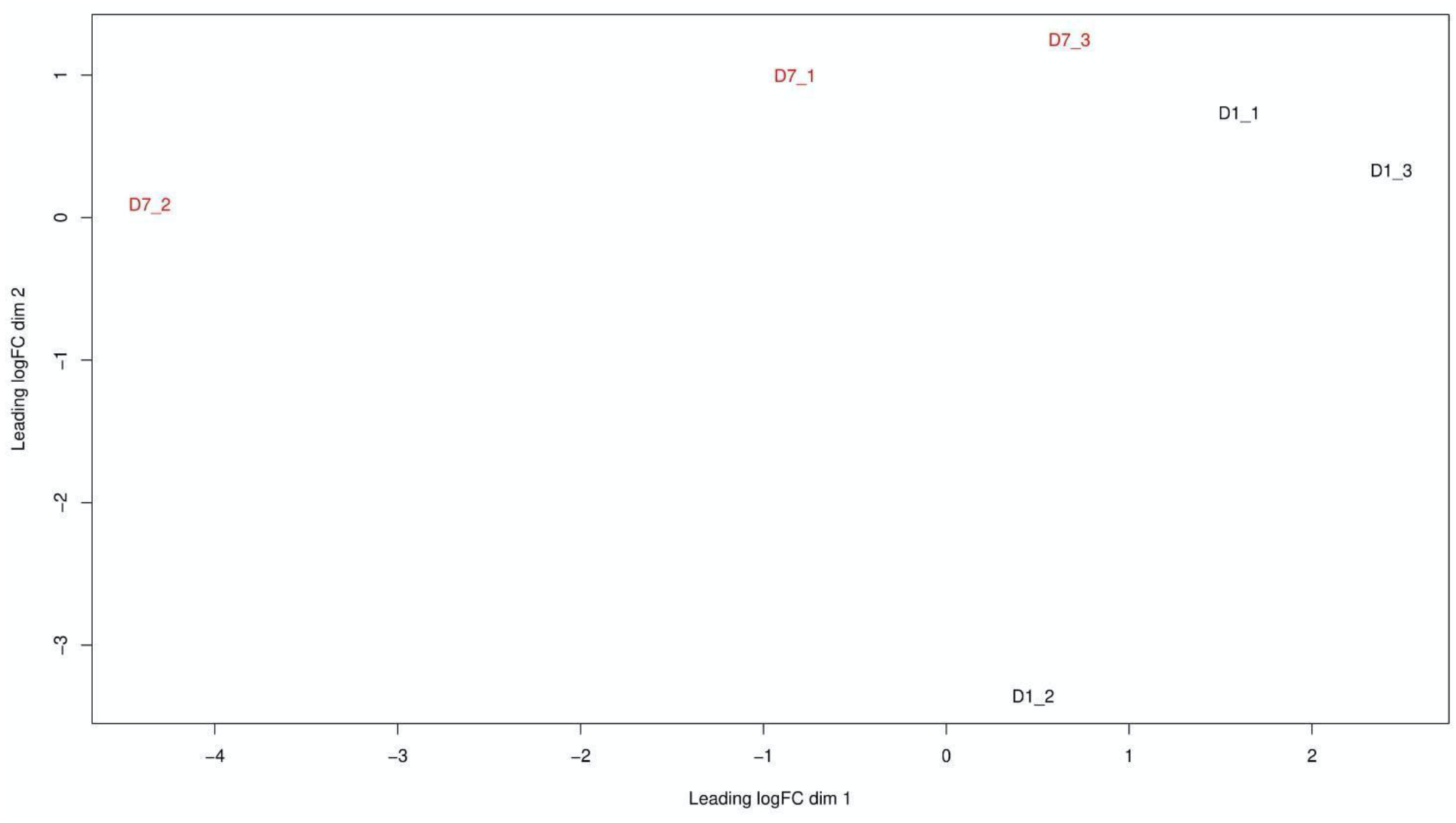
Multidimensional (MDS) Plot representing Day-1 and Day-7p.e. transcriptome datasets

Three biological replicates (1,2, and 3) of each Day-7 (in red) and Day-1 (in black) data sets were plotted in a Multidimensional plot to assess the similarity between them. The physical distance between the points is inversely proportional to their similarity.

