## Supplemental file3 for "Heterogeneous distribution of mRNAs within flight muscle fibers, and implications for function"

| **Significantly regulated genes within top 100 abundant genes averaged over Day1 and Day7** | **Base mean Value** | **Log_2_FC** | **Up or Down in Day1 (padj<0.05)** |
| --- | --- | --- | --- |
| sesB | 347850.7 | 2.227433 | UP |
| Mhc | 264281 | 5.709283 | UP |
| Mlc2 | 243145 | 4.524977 | UP |
| ATPsynC | 218363.7 | 3.354864 | UP |
| blw | 216409.7 | 3.324792 | UP |
| Act88F | 201461.6 | 7.238531 | UP |
| Gapdh1 | 148633.7 | 2.264797 | UP |
| Strn-Mlck | 126417 | 3.440295 | UP |
| kdn | 98516.44 | 3.624143 | UP |
| Eno | 86863.03 | 1.684224 | UP |
| fln | 76912.33 | 3.515013 | UP |
| CG9090 | 74523.35 | 2.347359 | UP |
| Mlc1 | 73166.5 | 2.786847 | UP |
| GstS1 | 61645.77 | 6.080123 | UP |
| up | 59463.3 | 3.353048 | UP |
| bt | 58783.95 | 1.86432 | UP |
| PyK | 57031.3 | 1.997068 | UP |
| UQCR-C1 | 56235.26 | 1.817812 | UP |
| Gpdh | 55297.21 | 3.06708 | UP |
| ATPsyngamma | 51655.79 | 2.122614 | UP |
| wupA | 50947.31 | 3.799695 | UP |
| l(1)G0156 | 49462.74 | 2.699792 | UP |
| Zasp52 | 48457.56 | 2.188995 | UP |
| Mdh2 | 47527.95 | 3.180395 | UP |
| GlyP | 44957.35 | 2.937401 | UP |
| Gpo-1 | 43347.7 | 2.34089 | UP |
| Unc-89 | 42479.78 | 3.146582 | UP |
| Tm1 | 40140.45 | 4.775185 | UP |
| Argk | 37275.35 | 2.666513 | UP |
| Tm2 | 36594.62 | 4.403918 | UP |
| Prm | 35667.59 | 1.517339 | UP |
| Pgi | 32387.49 | 2.071206 | UP |
| Gapdh2 | 32011.83 | 1.913473 | UP |
| Pfk | 26400.84 | 2.960475 | UP |
| muc | 29625.16 | 3.090952 | UP |
| CG7430 | 29097.91 | 2.345853 | UP |
| CG6439 | 25809.49 | 1.949463 | UP |
| CG5028 | 28742.38 | 1.618389 | UP |
| skap | 28324.28 | 1.952265 | UP |
| l(1)G0334 | 26619.82 | 2.17602 | UP |
| Pglym78 | 26611.7 | 2.608848 | UP |
| Mf | 174751.6 | -2.96652 | DOWN |
| mt:srRNA | 30692.35 | -2.12125 | DOWN |
| CG14645 | 28765.44 | -2.35999 | DOWN |
